# Butyrate reverts *Chlamydia pneumoniae* -induced transcriptional suppression of macrophage polarization markers and promotes chlamydial clearance in macrophages

**DOI:** 10.64898/2026.10.30.763131

**Authors:** Maarit Ylätalo, Inés Reigada, Leena Hanski

## Abstract

Short-chain fatty acids (SCFAs) are generated by gut microbiota through fermentation of dietary fibers. SCFAs contribute to human health through immunomodulation, metabolic regulation and other extraintestinal functions. The aim of this study was to investigate the effects of SCFAs on the survival and antibiotic susceptibility of the intracellular bacteria *Chlamydia pneumoniae* and *Chlamydia trachomatis*, both representing prevalent human pathogens associated with chronic sequelae. Despite earlier reports on their direct antimicrobial activity, none of the three major SCFAs acetate, propionate nor butyrate affected the growth of *C. pneumoniae* or *C. trachomatis* in epithelial cells. However, butyrate decreased *C. pneumoniae* genome copy numbers and infectious progeny production in THP-1 macrophages harbouring antibiotic-refractory infection phenotype. In addition, butyrate enhanced the chlamydiocidal efficacy of azithromycin in the macrophages. Similar results were obtained with *C. trachomatis*-infected macrophages. To elucidate the underlying molecular changes, impact of chlamydial infection, butyrate treatment and the combination theirof on macrophage polarization and antimicrobial peptide expression was investigated. *C. pneumoniae* suppressed the expression of macrophage polarization markers CD80, TLR4, CD64, CD204 and CD206, and the antimicrobial peptide LL-37, while butyrate treatment reverted the expression of both the polarization markers and LL-37 in the infected macrophages. These findings suggest that butyrate may play a role in modulating immune responses in macrophages during *Chlamydia* infections, highlighting the role of diet-derived metabolites in shaping *Chlamydia*–host interactions and antichlamydial treatment efficacy.

## 1 Introduction

The obligate intracellular *Chlamydia* species represent common bacterial infections causing agents globally. *Chlamydia pneumoniae* is a widespread respiratory pathogen, frequently implicated in community acquired pneumonias, sinusitis and bronchitis ^1^. It has also been associated with chronic obstructive pulmonary disease (COPD) and various stages of asthma ^2^. Beyond respiratory tract, *C. pneumoniae* has been isolated from atherosclerotic arteria and retina of Alzheimer’s disease patients, triggering research on the association of persistent chlamydial infection with chronic inflammatory diseases ^3^. Another chlamydial species, *C. trachomatis,* is the leading cause of sexually transmitted infections worldwide and associated with reproductive health complications such as ectopic pregnancies, infertility and pelvic inflammatory disease (PID). Furthermore, a subset of *C. trachomatis* serovars cause ocular infections and are recognized as the most common cause of infection induced blindness known as trachoma ^4^. *Chlamydia* spp. bacteria hence represent a significant public health burden both in high- and low-income countries.

The obligate intracellular lifestyle of *Chlamydia* is facilitated by a unique biphasic infection cycle, whose detailed dynamics and regulation remain yet to be understood ^5, 6^. In a productive infection, which occurs for example in lung or genital mucosa epithelium, *Chlamydia* switches between an infectious form, elementary bodies (EBs) and intracellular, replicative form reticulate bodies (RBs). Various experimental models have demonstrated the existence of a third morphological form of *Chlamydia,* known as aberrant bodies (ABs) ^7, 8^. ABs are characterized as enlarged, non-replicative and metabolically slow forms of bacteria which arise in response to antibiotics, nutrient deprivation or other external stress factors. Besides this specific morphology, the chlamydial stress response, often referred as chlamydial persistence, is associated with a characteristic gene expression profile, a viable but non-cultivable phenotype and compromised antibiotic susceptibility. The persistent state is considered reversible, allowing the bacterium to return to its productive developmental cycle and continue replication when stressful conditions clear out. Even though definitive evidence on the causal relationship of persistence in clinical chlamydial infections to their contribution to chronic inflammatory conditions *in vivo* remains incomplete, the antibiotic-refractory nature and recurrent nature of *Chlamydia* in the host are commonly associated with persistence. Innate immune cells, particularly monocytes and macrophages, play a key role in this respect. When internalized into these cells, *Chlamydia* exhibit the hallmarks of above-mentioned stress response, including compromised eradication by antibiotics ^9, 10^. Azithromycin, the first-choice antibiotic for chlamydial infections, has a minimum inhibitory concentration MIC of 25 nM against *C. pneumoniae* in epithelial cells ^7^, whereas in macrophages concentrations in the range of 1 µM are required for eradication ^11^. Furthermore, infected myeloid cells are thought to mediate chlamydial dissemination from the primary site of infection to local lymph nodes and other body sites.

Macrophages have a central role in innate immune responses against pathogens, but they are also key contributors to inflammation and its resolution in various non-infectious pathologies. Macrophages represent physiologically and phenotypically diverse and plastic cell populations which classically have been thought to adopt two different polarization directions, M1 or M2 ^12^. M1, so-called classically activated macrophages, are proinflammatory, promoting immune responses to fight infections. They produce proinflammatory cytokines and microbicidal reactive oxygen and nitrogen species. M2, or alternatively activated macrophages, are anti-inflammatory and involved in tissue repair and inflammation resolution. Functional phenotypes can alter between M1 and M2, and M1 and M2 transcriptional programs can also interfere with each other, leading to mixed gene expression profiles and function ^13^.

As intracellular pathogen *Chlamydia* have evolved strategies to manipulate macrophage function to avoid the host immune response ^14, 15^. *C. pneumoniae* has been described to effect macrophage polarization during the infection ^16^ but the effect appears cell type- and species-specific ^8^. Earlier research also suggests that pre-polarized M1 or M2 macrophages influence chlamydial replication, persistence and survival differently ^8, 17, 18^.

Extensive body of research has established the central role of intestinal microbiota in human health ^19^. Besides its direct impact on gut physiology, gut microbiota can have a significant effect on extraintestinal functions, such as respiratory tract physiology and lung diseases ^20^. The crosstalk between the intestinal and pulmonary systems, commonly referred as the gut-lung axes, involves the cell responses, host and microbe signaling pathways and the action of bacterial metabolites, such as short chain fatty acids (SCFAs). SCFAs are small carboxylic acids produced mainly in the large intestine by the gut microbiota through fermentation of dietary fibers. Among the different SCFAs, butyrate in particular has been reported to alleviate pulmonary diseases, such as asthma and COPD through various immunomodulatory effects ^21^. Butyrate levels have been reported to negatively correlate with symptom severity in obstructive lung disease patients ^22^ and with allergic and non-allergic asthma ^23^. Intestinal absorption of SCFAs results in micro- or millimolar concentrations in the lungs among other organs ^24^. In addition to gut microbiota, SCFAs are produced by vaginal microbiota, mainly during bacterial vaginosis ^25^.

Earlier research suggests that at micromolar concentrations, butyrate can exert anti-inflammatory actions by suppressing cytokine production and promoting alternative macrophage activation route ^26^. However, other reports indicate that millimolar butyrate may enhance pro-inflammatory and antimicrobial effector functions, including inflammasome activation and nitric oxide production ^27, 28^. While the varying butyrate concentrations applied in these studies hinder direct comparisons, controversies in previous reports presumably reflect the heterogeneity of macrophage populations and experimental models.

Butyrate’s ability to modulate the expression of immunomodulatory genes occurs by the inhibition of histone deacetylase (HDAC) activity ^29, 30^. HDAC acts by removing acetyl groups from histones, and suppression of HDAC activity leads to more open chromatin and accessible genes and thus induced transcription. Previous research connects HDAC inhibition to macrophage phenotypes ^31, 32^, increased inflammasome activity ^33^, and antimicrobial functions such as increased autophagy ^28^.

Despite the well-established impact of butyrate on macrophage physiology, there is currently no information about the impact of butyrate or other SCFAs on chlamydial infection. In this work, we studied the impact of SCFAs on *C. pneumoniae* and *C. trachomatis* infection in both epithelial cells and THP-1 macrophages. Since previous research links macrophage polarization status to chlamydial replication and persistence, we also wanted to elaborate whether THP-1 macrophage polarization is altered during *C. pneumoniae* infection and butyrate treatment. Drawing on previously reported findings on the ability of butyrate to induce the expression of the antimicrobial peptide LL-37, we also investigated the connection of LL-37 expression to the antichlamydial effects of butyrate.

## 2 Materials and Methods

### 2.1 Dosage information

Sodium acetate, sodium propionate and sodium butyrate were obtained from Sigma-Aldrich (St. Louis, Missouri, USA, purity-% ≥ 98). Stock solutions (20 mM) were prepared in sterile H_2_O and sterilized by filtration through 0.2 µm filter. Cells were exposed to SCFAs at final concentrations of 100 and 200 µM, selected to represent physiologically relevant exposure levels ^24^ and commonly used concentrations in cell culture studies ^34^ . Cell viability data is available in Supplementary information 1.

### 2.2 Cell culture

A549 cells (ATCC CCL-185) were maintained in low glucose DMEM medium (Biowest, Nuaillé, France), supplemented with 10% FBS (Biowest), 2 mM L-gutamine (Sigma-Aldrich) and 20 µg/ml gentamicin (Gibco, Thermo Fisher Scientific, Grand Island, NY, USA). Before experiments, cells were seeded into well plates at indicated densities and allowed to adhere overnight.

HeLa 229 cells (ATCC CCL-2.1) were maintained in high glucose DMEM medium (Cytiva, Logan, UT, USA), supplemented with 10% FBS, 2 mM L-gutamine, 1 mM sodium pyruvate (Sigma-Aldrich) and 20 µg/ml gentamicin. Before experiments cells were seeded into well plates at indicated densities and allowed to adhere overnight.

THP-1 (ATCC TIB-202) cells were maintained in RPMI-1640 medium (Cytiva), supplemented with 10% FBS, 0.05 mM 2-mercaptoethanol (Gibco) and 20 µg/ml gentamicin. To differentiate THP-1 cells to macrophages, cells were seeded into well plates at indicated densities and treated with 160 nM phorbol-12-myristate-13-acetate (PMA, Sigma-Aldrich) for 72 h.

### 2.3 Bacterial strains and infections

*C. pneumoniae* cardiovascular isolate CV6 was obtained from Professor Matthias Maass (Paracelsus Medical University, Salzburg, Austria) and propagated as previously described in ^35^. *C. trachomatis*, serovar K (ATCC UR-887), obtained from ATCC, was propagated in HeLa 229 cells in high glucose DMEM medium, supplemented with 10% FBS, 2 mM L-glutamine, 1 mM sodium pyruvate, 20 µg/ml gentamicin, 0,6 µg/ml cycloheximide, 100 µg/ml streptomycin and 3.75 µg/ml amphotericin B. Cells were inoculated with *C. trachomatis* and incubated 48 h. After that, cells were lysed, debris was removed by centrifugation (10 min, 500 g) and *C. trachomatis* EBs were collected by pelleting (1 h, 21 000 g, 4 °C). Both chlamydial strains were stored at −80 °C in sucrose-phosphate-glutamic acid (SPG) ^36^. Cells were infected with multiplicity of infection (MOI) 1 in all experiments. Inoculum was added to the cells, and the plates were centrifugated at 550 g for 1 h and incubated at 37 °C another hour. Then, inoculum was removed and cell culture medium with possible study compounds was added. 1 µg/ml of cycloheximide (CHX) was added to the cells in experiments with A549 and HeLa 229 cells to maintain stable host cell numbers throughout the infection.

### 2.4 Determining *Chlamydia* growth and Infectious progeny formation

A549 and HeLa 229 cells were seeded into 24-well plate, on top of coverslips, at a density of 4×10^5^ cells/well, infected with *C. pneumoniae* or *C. trachomatis*, respectively, at MOI1 as described above. At 72 h (*C. pneumoniae*) or 48 h (*C. trachomatis*) post-infection, cells were fixed with methanol for 10 minutes and blocked with 0,5 % BSA. Chlamydial inclusions were then stained with Chlamydia LPS Monoclonal antibody (B410F, Invitrogen, Thermo Fisher Scientific, 1:100) for 1 h. After washing 3 times with PBS, Alexa Fluor^TM^ 488 (Invitrogen, Thermo Fisher Scientific 1:2000) was used as the secondary antibody. Inclusions were quantified with fluorescence microscopy. In infectious progeny formation assay THP-1 (3.5×10^5^ cells/well), A549 (4×10^5^ cells/well) or HeLa (4×10^5^ cells/well) cells were infected with MOI1 and treated as described above. At 48 or 72 h post infection, newly formed EBs were harvested by scraping the cells from the wells, lysed, and fresh A549 or HeLa cells were infected with the collected lysates. After infection, inoculum was removed, and cells were incubated for 48 or 72 h with fresh cell culture medium containing 1 µg/ml of CHX. Chlamydial inclusions were stained as described above and inclusions quantified with fluorescent microscopy.

### 2.5 Quantitative PCR

To determine *Chlamydia* genome copy numbers, A549, HeLa 229 or THP-1 cells were seeded into 24-well plate as described above and infected with *C. pneumoniae* or with *C. trachomatis* and treated with SCFAs. At 48 or 72 h post-infection, cells were collected, lysed and DNA was extracted with GeneJet Genomic DNA purification kit (Thermo Fisher Scientific) according to the manufacturer’s instructions for mammalian cells. Chlamydial genome copy numbers were determined with Step One plus Real-Time PCR system (Thermo Fisher Scientific). The primers used for *C. pneumoniae* were against OmpA gene (forward primer: 5′-TCC GCA TTG CTC AGC C-3′ and reverse primer 5′-AAA CAA TTT GCA TGA AGT CTG AGA A-3′) ^37^ and the primers for *C. trachomatis* were against 16S RNA gene (forward primer: 5’-GGC GTA TTT GGG CAT CCG AGT AAC G-3’ and reverse primer: 5’-TCA AAT CCA GCG GGT ATT AAC CGC CT-3’) ^38^.

### 2.6 Reverse transcription PCR

To determine the gene expression of M1 and M2 macrophage markers, RT-PCR was conducted as described in Ylätalo et al 2025 ^39^. Briefly, THP-1 cells were seeded into 24-well plate as described above, differentiated and infected with *C. pneumoniae* and treated with 200 µM butyrate. After 72 h, cells were lysed, and RNA was extracted with PureLink RNA Mini Kit (Thermo Fisher Scientific). cDNA synthesis was conducted with RevertAid First Strand cDNA Synthesis Kit (Thermo Fisher Scientific), and the expression of studied genes was determined with Step One Plus Real-Time PCR system (Thermo Fisher Scientific), using GAPDH as an endogenous reference gene. Primers used in this study are listed in Supplementary Table 1.

### 2.8 Data analysis

Statistical analyses were performed using IBM SPSS Statistics version 30. Data are presented as mean ± SEM of at least three biological replicates. Differences between the groups were analyzed by One-way ANOVA followed by Games-Howell post hoc test.

Statistical significances are presented as *p* values, with *p* 0.05 considered statistically significant.

In gene expression analysis, relative gene expression levels were normalized to the housekeeping gene GAPDH and calculated using 2^-ΔΔCt method and presented as relative quantification (RQ) values.

## 3 Results

### 3.1 Impact of SCFAs on *C. pneumoniae* infection in epithelial cells

To study the impact of acetate, propionate and butyrate on *C. pneumoniae* growth and infectious progeny formation, A549 lung epithelial cells were infected with MOI1 of *C. pneumoniae* and treated with 100 µM and 200 µM of sodium acetate, sodium propionate and sodium butyrate. As illustrated in figure 1, none of the SCFAs had an effect on *C. pneumoniae* growth as determined by inclusion counts at 72 h post-infection (Fig. 1A) or infectious progeny production (Fig 1B).

**Figure 1.**
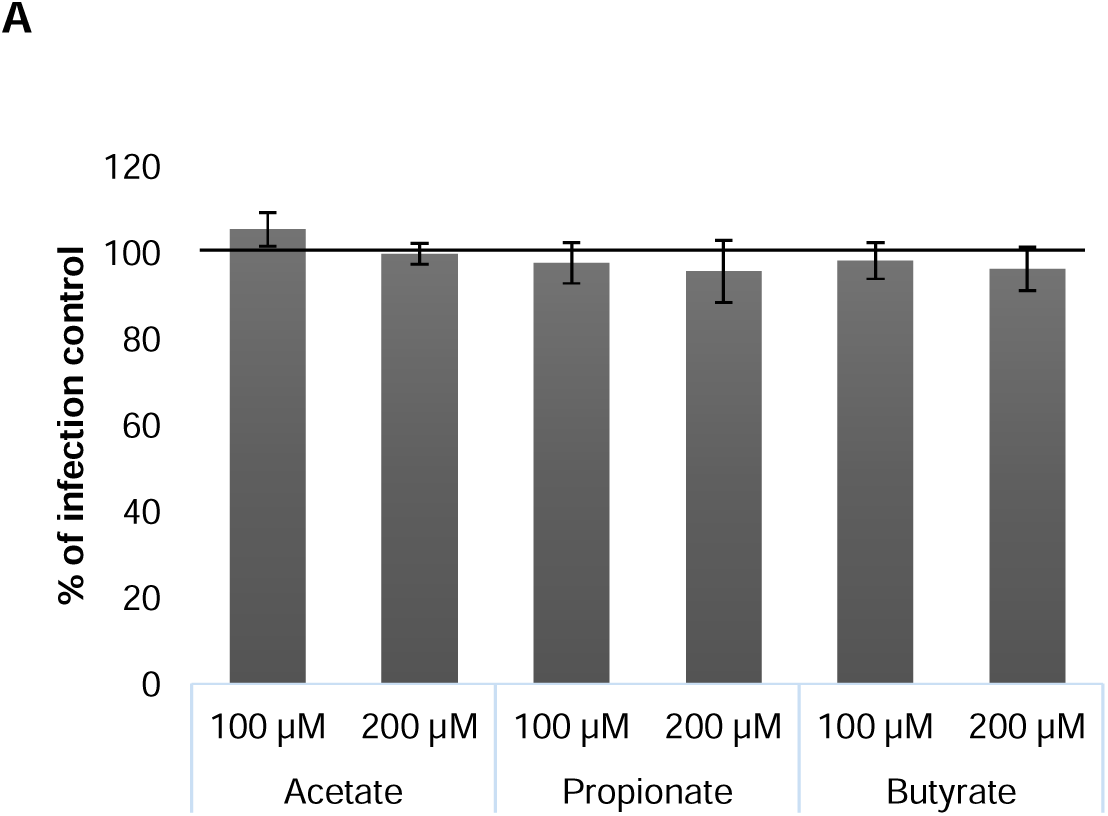
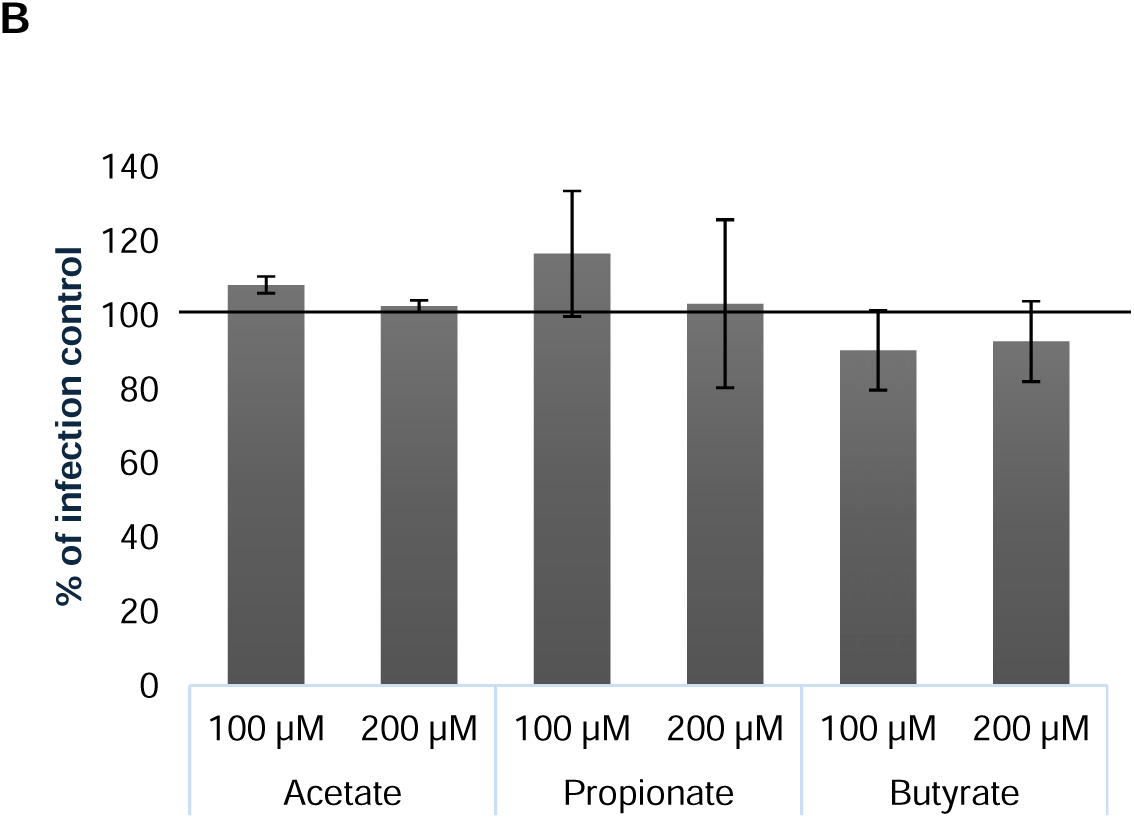
Impact of SCFAs on *C. pneumoniae* infection in A549 epithelial cells. A) *C. pneumoniae* inclusion counts and B) infectious progeny production after 72 h treatment with the SCFAs. Results are normalized to nontreated infection control and presented as mean ± SEM. Statistical significances were calculated using ANOVA and Games-Howell post hoc test. n ≥ 4.

### 3.2 Impact of SCFAs on *C. pneumoniae* infection in THP-1 macrophages

To determine the impact of SCFAs on *C. pneumoniae* infection in THP-1 macrophages, a chlamydial infectious progeny assay was performed.

As shown in Figure 2A, acetate and propionate had no effect on *C. pneumoniae* infectious progeny production in THP-1 macrophages, whereas butyrate decreased the *C. pneumoniae* infectious progeny yield statistically significantly, by 60% (*p* = 0.028).

**Figure 2.**
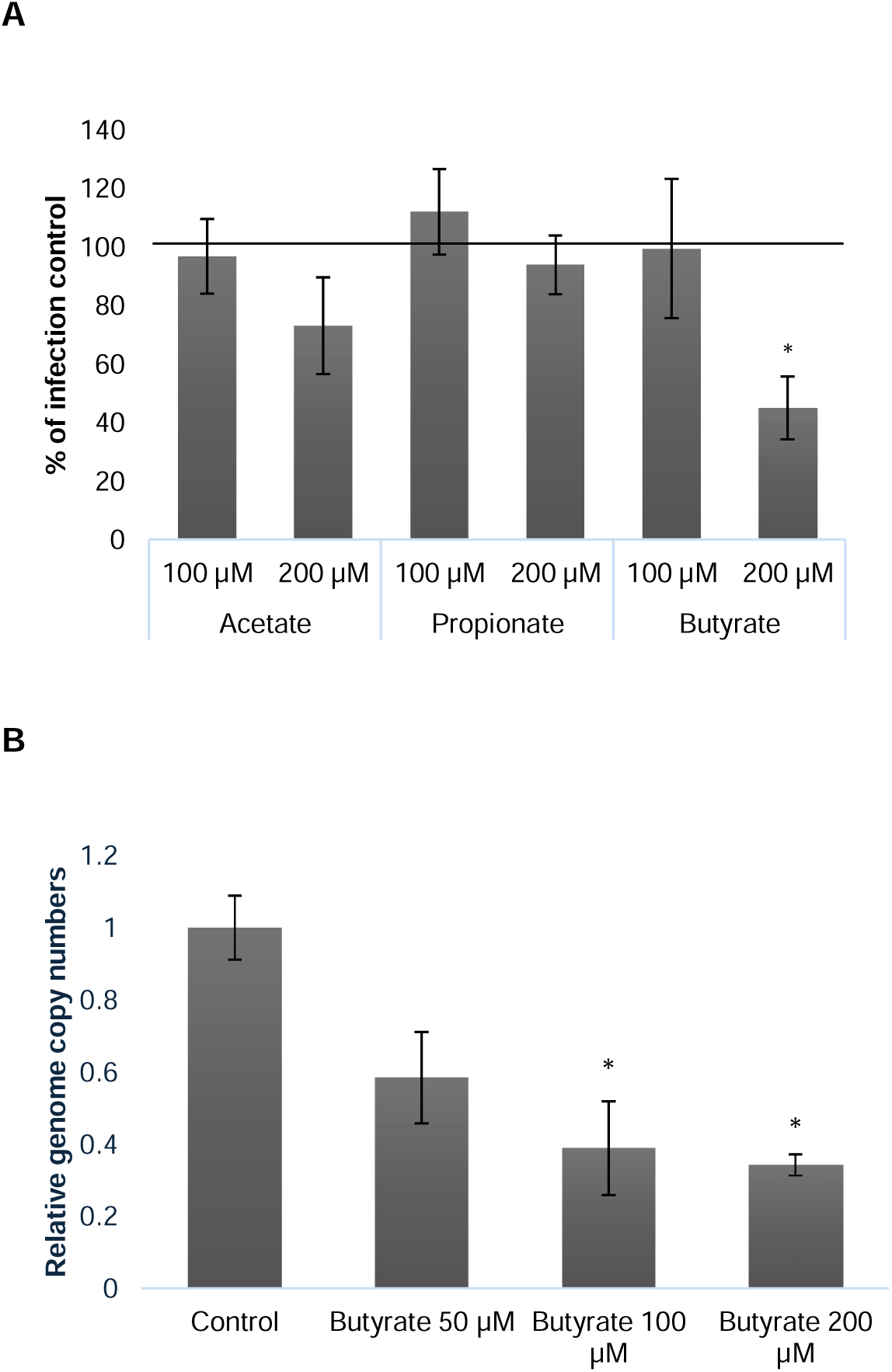
Impact of SCFAs on *C. pneumoniae* infection in THP-1 macrophages. A) infectious progeny production after 72 h treatment with the SCFAs; B) genome copy numbers after 72 h treatment with different concentrations of butyrate. Results are normalized to nontreated infection control and presented as mean ± SEM. Statistical significances were calculated using ANOVA and Games-Howell post hoc test (*p* < 0.05: _∗_), n ≥ 4.

As the infectious progeny quantification data indicated that butyrate may suppress *C.* pneumoniae growth in the macrophages, we subsequently confirmed this effect by detecting *C. pneumoniae* DNA replication in the cells with quantitative PCR. As shown in figure 2B, butyrate at concentrations 100 µM and 200 µM decreased *C. pneumoniae* genome copy numbers statistically significantly, 100 µM by 61 % (*p* = 0.038) and 200 µM by 66 % (*p* = 0.01).

### 3.3 Impact of SCFAs on the efficacy of azithromycin on *C. pneumoniae* infection in THP-1 macrophages

Since the standard antibiotic used for chlamydial infections, azithromycin, requires much higher concentrations to eradicate the infection in macrophages than in epithelial cells, we next examined whether SCFA treatment enhances its antichlamydial efficacy in macrophages. To this end, *C. pneumoniae* infectious progeny and genome copy number (qPCR) analyses were performed to compare the effect of butyrate (200 µM), azithromycin (100 nM) alone and as combination. Butyrate improved azithromycin’s effect by decreasing *C. pneumoniae* infectious progeny production statistically significantly by 78 % (*p* = 0.038) compared to only azithromycin (Fig. 3A). According to qPCR data, butyrate did not affect the capacity of azithromycin to reduce *C. pneumoniae* genome copy numbers in the macrophages (Fig. 3B). Azithromycin and butyrate monotherapy showed similar impact on *C. pneumoniae* genome counts by decreasing them by 68 % and 58 %, respectively, while the combination decreased the genome copy numbers by 68 %. Collectively, these results imply that butyrate improves the chlamydiocidal activity of azithromycin on *C. pneumoniae,* yet residual chlamydial DNA levels are not affected compared to azithromycin monotherapy. In qPCR analysis, DNA of dead bacteria that yet haven’t been degraded by the host, are present in the samples, which might influence the outcome of this experimental setup.

**Figure 3.**
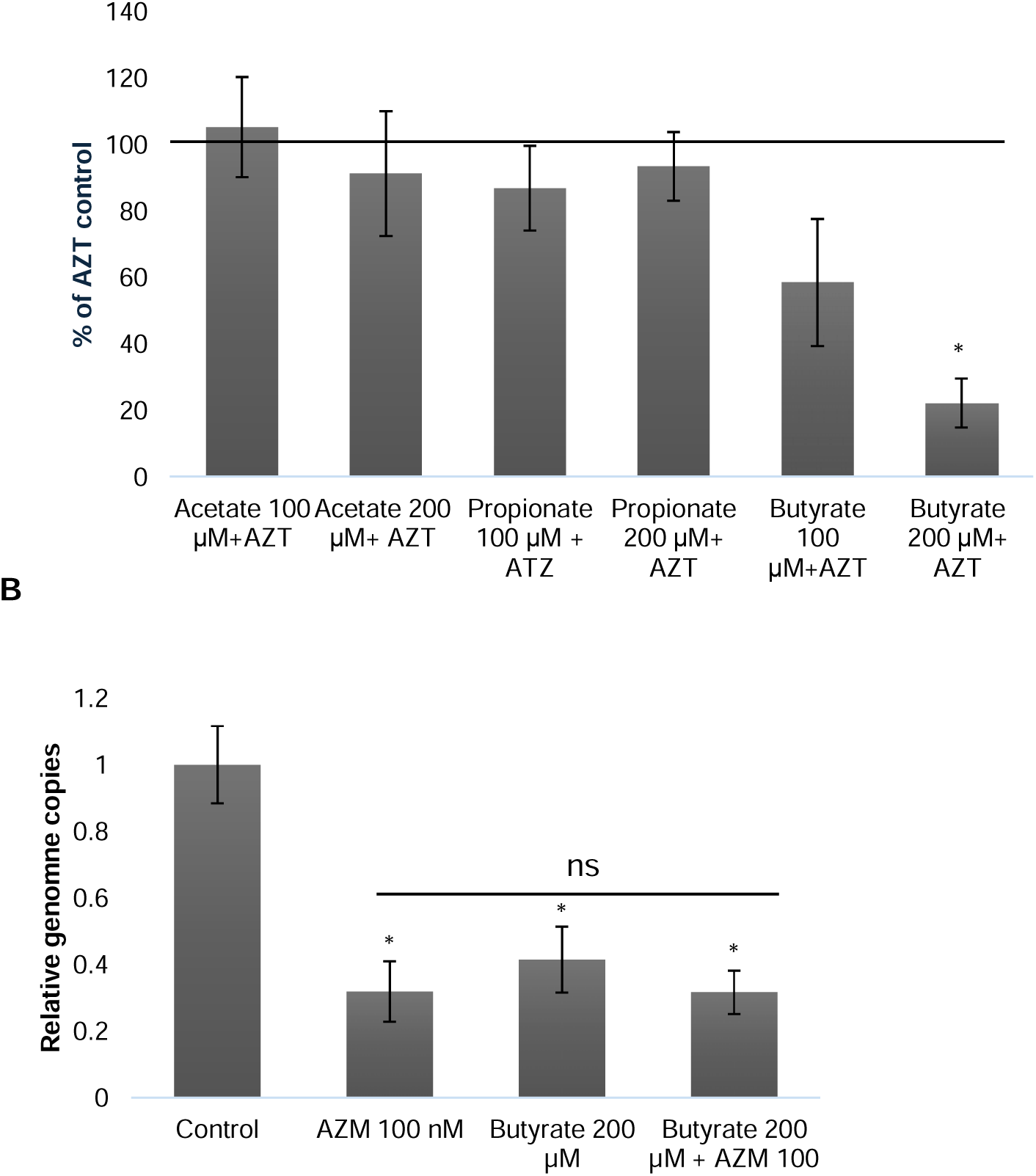
Impact of SCFAs on the efficacy of azithromycin (AZM) against *C. pneumoniae* in THP-1 macrophages. A) Infectious progeny production and B) genome copy numbers of *C. pneumoniae* at 72 h post-infection, treated with butyrate, azithromycin or combination. Results are normalized to azithromycin-treated infection control and presented as mean ± SEM. Statistical significances were calculated using ANOVA and Games-Howell post hoc test (*p* < 0.05: _∗_; < 0.01: _∗∗_; < 0.001: _∗∗∗_). n = 4.

### 3.4 Impact of butyrate on *C. trachomatis* infected HeLa epithelial cells and THP-1 macrophages

To study whether butyrate exerts similar effects also on other *Chlamydia* species, HeLa-229 cells were infected with *C. trachomatis*, MOI1, and treated with 200 µM butyrate, the effective concentration against *C. pneumoniae*. Bacterial growth and infectious progeny formation of *C. trachomatis* were determined. Consistent with the data obtained on *C. pneumoniae* infections in A549 cells, there were no differences between nontreated infection control and butyrate treated samples either in the chlamydial inclusion counts or infectious progeny formation (Fig. 4). These data imply that butyrate has no effect on *Chlamydia* infection in permissive cells, such as epithelial cells, where most of the cells are in actively replicative form. Azithromycin is very effective against actively replicating *C. trachomatis* in epithelial cells. The same concentration that fails to eliminate the infection from macrophages results practically 100% eradication in epithelial cells (data not shown).

**Figure 4.**
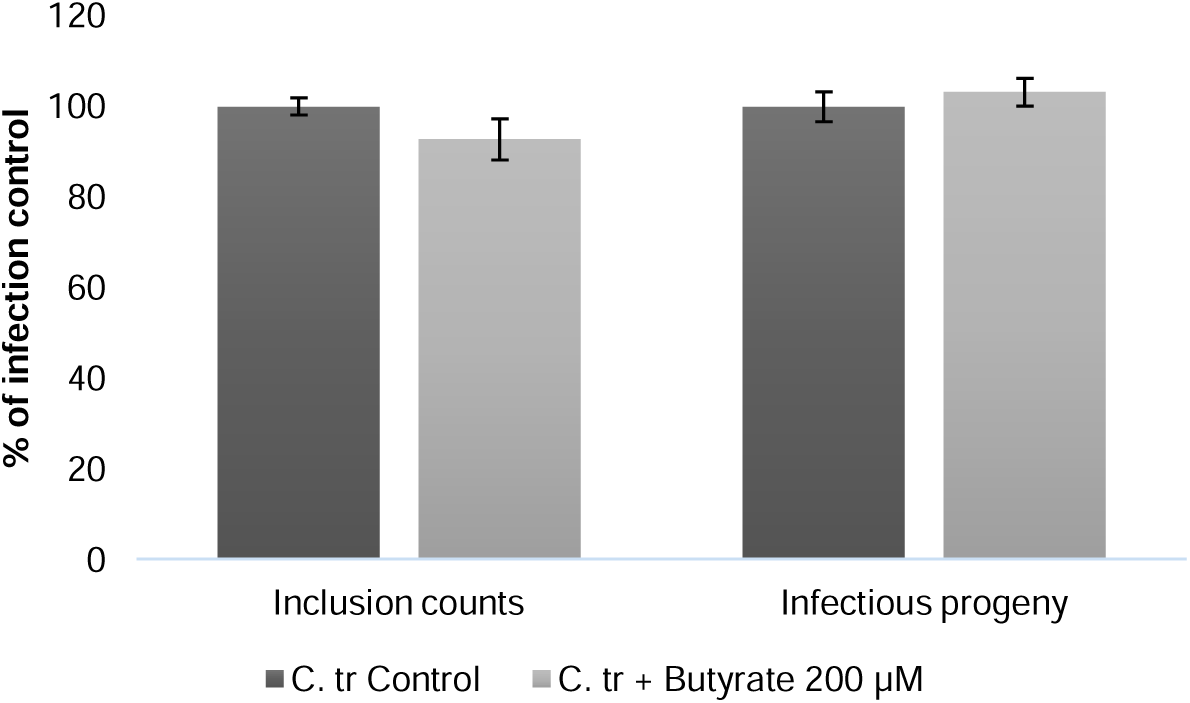
Impact of butyrate on *C. trachomatis* infection in HeLa-229 cells. *C. trachomatis* inclusion counts and infectious progeny production at 48 h post-infection. Results are normalized to nontreated infection control and presented as mean ± SEM. Statistical significances were calculated using ANOVA and Games-Howell post hoc test. n ≥ 3.

Next, the impact of butyrate on *C. trachomatis* infection and the efficacy of azithromycin in THP-1 cells were determined. THP-1 macrophages were infected and treated with butyrate with or without azithromycin. Butyrate decreased the *C. trachomatis* genome copy numbers by 23 % (*p* = 0.012) and azithromycin by 65 % (*p* = 0.0002), shown in Figure 5A. As shown in Fig. 5B, our results indicate that butyrate seemed to decrease also the *C. trachomatis* infectious progeny formation by 29.8 %, but it did not reach statistical significance, as did azithromycin by decreasing the infectious yield by 64.9 % (*p* = 0.007). Combination of azithromycin and butyrate was even more effective and decreased the infectious progeny formation by 93,7 % (p 0.001), compared to infected control.

**Figure 5.**
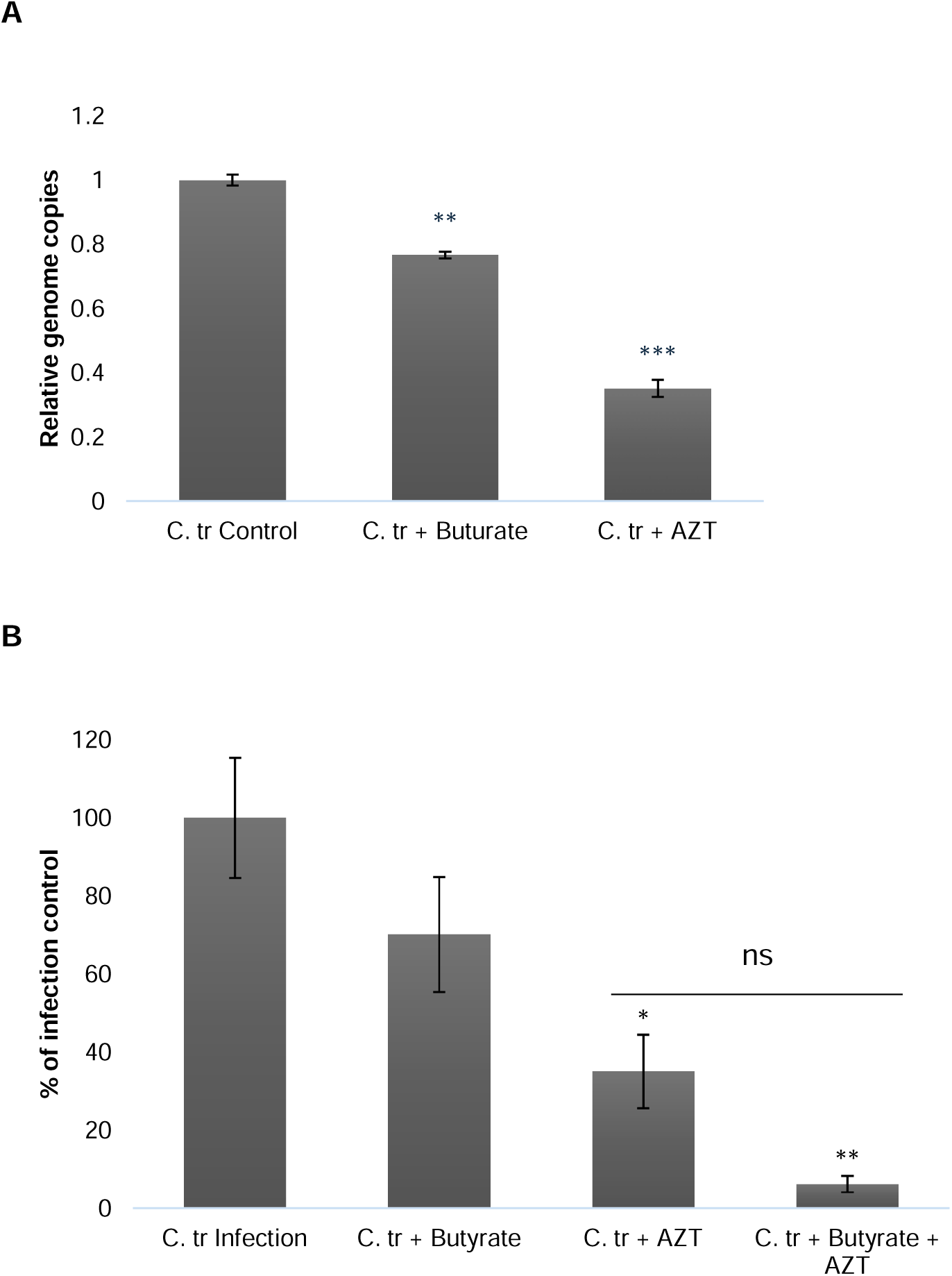
Impact of butyrate (200 µM) on *C. trachomatis* infection and its susceptibility to azithromycin (AZT, 100 nM) in THP-1 cells. A) Genome copy numbers and B) Infectious progeny production at 48 h post-infection, treated with butyrate, azithromycin and combination. Results are normalized to nontreated infection control and presented as mean ± SEM. Statistical significances were calculated using ANOVA and Games-Howell post hoc test (*p* < 0.05: _∗_; < 0.01: _∗∗_; < 0.001: _∗∗∗_), n ≥ 3.

### 3.5 Impact of *C. pneumoniae* and butyrate on THP-1 macrophage polarization

Previous research indicates that *C. pneumoniae* infection may alter macrophage polarization, which can contribute to their immune evasion. It is also known that butyrate affects macrophage polarization, yet this seems to be cell type dependent. Hence it was of interest to evaluate the impact of *C. pneumoniae* infection, butyrate treatment and their combination on THP-1 macrophages to gain insights on its potential connection to the macrophage-specific antichlamydial effects observed with butyrate. Macrophage M1 and M2 polarization marker expression were evaluated by quantifying the mRNA levels. The results of this analysis are shown in Figure 6. CD80, TLR4 and CD64 are classic M1 polarization markers and CD204 and CD206 are M2 polarization markers ^40, 41^. According to our data, *C. pneumoniae* infection suppressed expression of all markers, RQ-values being CD80 = 0.80, TLR4 = 0.50, CD64 = 0.35, CD204 = 0.50 and CD206 = 0.51. Treating non-infected macrophages with butyrate had only a slight effect on the expression of the markers (RQ values: CD80 = 1.2, TLR4 = 0.92, CD64 = 1.24, CD204 = 1.35 and CD206 = 1.23).

**Figure 6.**
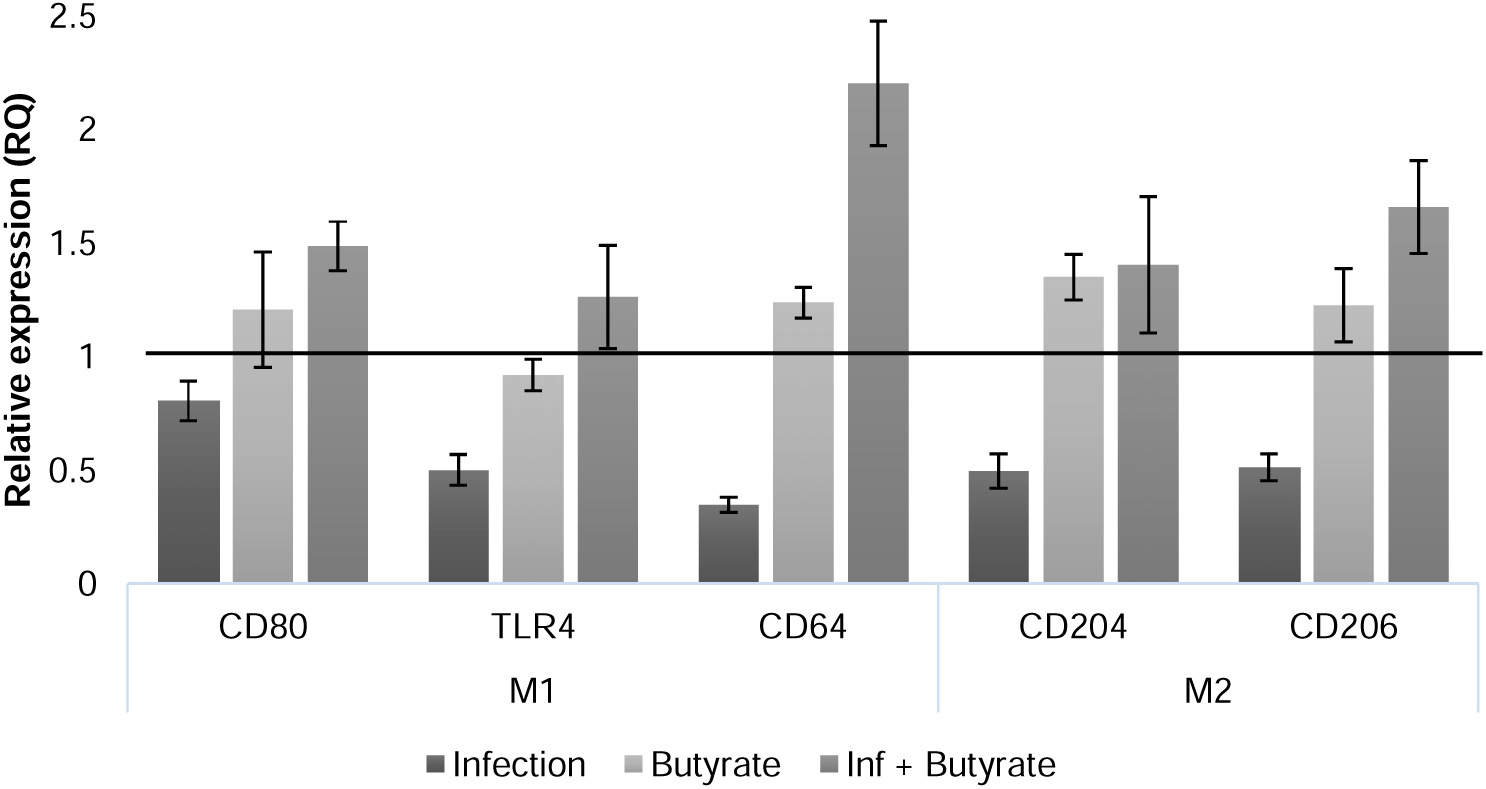
Impact of *C. pneumoniae* infection and butyrate on THP-1 macrophage polarization. Expression of the markers was determined at 72 h post-infection by comparing the amplification of each gene compared to the internal reference gene GAPDH. Infection samples and non-infected butyrate-treated samples were compared to untreated control, while Infection + butyrate was compared to infected samples. Relative quantification (RQ) values were obtained using 2^-ΔΔCt method. RQ values (±SEM) are presented. n = 6.

However, in *C. pneumoniae* -infected macrophages butyrate treatment increased the expression of the markers. Compared to infected control, CD64 marker was the most upregulated (RQ 2,2), while CD80 (RQ = 1.49), TLR4 (RQ = 1.26), CD204 (RQ = 1.40) and CD206 (RQ = 1.66) were more moderately elevated. These results indicate that butyrate may overwrite *C. pneumoniae* induced suppression of the macrophage activation and induce the polarization of infected cells.

### 3.6 Induction of the LL-37 antimicrobial peptide and its role in *C. pneumoniae* infection in THP-1 macrophages

Earlier research indicates that butyrate upregulates the expression of the antimicrobial peptide LL-37 in various cell lines and that LL-37 has antimicrobial activity against several pathogens ^42^. To study the relevance of LL-37 in the macrophage-specific antichlamydial effect of butyrate, we quantified the expression of LL-37 in *C. pneumoniae* -infected and butyrate-treated macrophages by RT-PCR. According to our results, butyrate did not alter LL-37 expression in non-infected THP-1 macrophages (RQ = 1.1; Fig. 7A). In contrast, our data indicate that *C. pneumoniae* infection downregulates LL-37 expression (RQ = 0,41) and when the infected macrophages were treated with butyrate, the LL-37 expression was recovered and even upregulated (RQ = 1,78).

**Figure 7.**
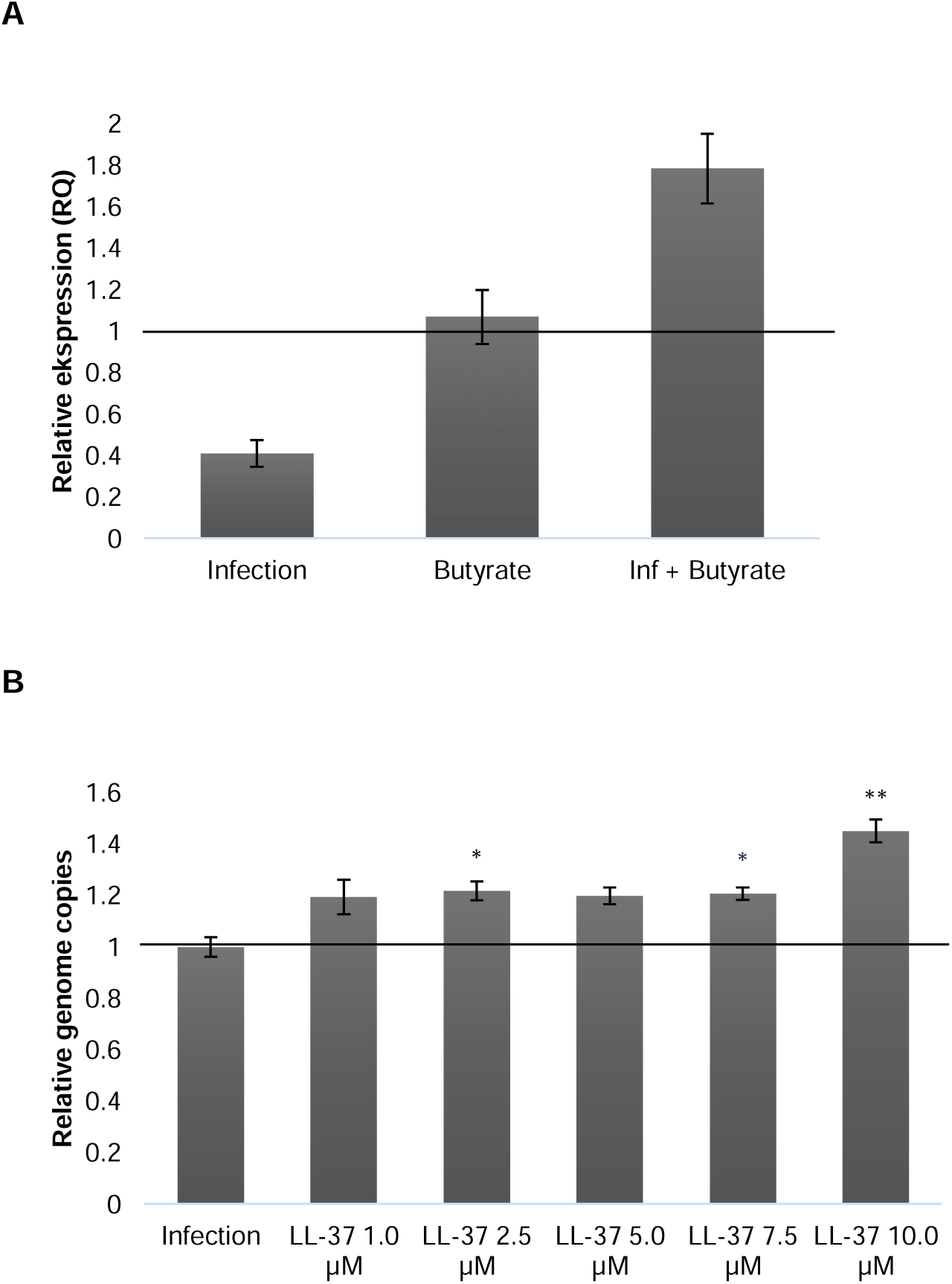
Role of the antimicrobial peptide LL-37 in butyrate-induced changes in THP-1 macrophages. A) Impact of butyrate, *C. pneumoniae* infection and their combination on LL-37 expression. RNA was collected at 72 h post-infection, and the amplification of LL-37 gene was compared to the internal reference gene GAPDH. Relative quantification (RQ) values were obtained using 2^-ΔΔCt method. RQ values (±SEM) are presented. n = 6. B) Impact of LL-37 supplementation on *C. pneumoniae* genome copy numbers at 72 h post-infection. Results are normalized to nontreated infection control and presented as mean ± SEM. Statistical significances were calculated using ANOVA and Games-Howell post hoc test (*p* < 0.05: _∗_; < 0.01: _∗∗_; < 0.001: _∗∗∗_). n = 4.

To evaluate the role of LL-37 expression recovery in butyrate’s ability to suppress *C. pneumoniae* infection in THP-1 macrophages, we studied the impact of externally added LL-37 on *C. pneumoniae* infected macrophages. When infected cells were treated with LL-37 in various concentrations (1.0, 2.5, 5.0, 7.5 and 10.0 µM), none of them decreased *C. pneumoniae* genome copy numbers quantified at 72 h post-infection (Fig 7B). In fact, all studied concentrations of LL-37 increased the *C. pneumoniae* genome copy numbers (1,0 µM by 19,5 %, 2,5 µM by 21.8 %, 5.0 µM by 19.9 %, 7.5 µM by 20.1 % and 10.0 µM by 45.1%).

## 4 Discussion

In clinical practice, *Chlamydia* infections are usually treated with azithromycin, other macrolides or doxycycline ^43, 44^, However, although several antibiotics show low MIC values in epithelial cells, conventionally used for antimicrobial susceptibility testing, eradication of bacteria from macrophages requires substantially higher concentrations ^10, 11, 45^. This illustrates the importance of macrophages and other myeloid cells in chlamydial infections and calls for a better understanding on physiological and dietary factors governing antichlamydial treatment efficacy in these cells.

In this study, we demonstrate that butyrate inhibits *C. pneumoniae* and *C. trachomatis* infection in THP-1 macrophages (Fig. 2 and 5, respectively) where these bacteria exhibit a persistent phenotype, while having no effect on chlamydial growth in epithelial cells (Fig. 1 and 4). These findings reveal cell type-specific effects of butyrate and suggest that its antichlamydial activity is mediated through modulation of host immune responses rather than direct antichlamydial effects. Consistent with our earlier observations ^10, 11^, azithromycin failed to eradicate *Chlamydia* from THP-1 macrophages (Fig 3 and 5), while combined treatment with butyrate enhanced its efficacy. Concomitant treatment decreased the *C. pneumoniae* infectious progeny production statistically significantly compared with azithromycin alone (Fig. 3A). The lack of differences in *C. pneumoniae* genome copy numbers between azithromycin and azithromycin + butyrate treatments (Fig. 3B) imply that butyrate can enhance the azithromycin’s antichlamydial activity by affecting bacterial infectivity rather than overall bacterial load. A similar trend was observed with *C. trachomatis,* but the results did not reach statistical significance (Fig. 5).

Butyrate’s ability to modulate macrophage polarization has been addressed in various earlier studies reporting either promotion of M2 polarization ^26, 46, 47^, suppression of M2 markers or downregulated M1 activation ^48^. In THP-1 macrophages, exposed to LPS, butyrate induces caspase 1-activation and NO release, which are classical M1 macrophage functions ^27^, whereas another study reported inhibition of M1 markers, including CD80 ^34^. Collectively, these findings indicate that immunomodulatory effects of butyrate are highly dependent on the cellular context and activation environment.

Here, we determined the expression of classical macrophage M1 and M2 polarization markers in *C. pneumoniae* -infected and/or butyrate-treated THP-1 macrophages. CD80, CD64 and TLR4 were selected as M1 markers ^41, 49, 50^ and CD204 and CD206, as M2 markers ^50, 51^. *C. pneumoniae* infection suppressed all the studied markers (Fig. 6), implying a tolerized, dysregulated or hypoactive macrophage phenotype ^52, 53^. Such phenotype may benefit the pathogen by limiting both microbial killing and inflammation resolution ^54^.

Immunosuppressed macrophages may provide a permissive intracellular niche, facilitating chlamydial survival. This observation aligns with earlier reports that certain intracellular pathogens, such as *Salmonella, Mycobacteria* and *Coxiella* actively modulate macrophage polarization to evade host immune responses and persist in the cells ^54–56^. While the above-mentioned studies have reported intracellular bacteria mostly shift macrophages toward anti-inflammatory polarization states, our data explicitly show a broad suppression across classical M1 and M2 markers in the context of *C. pneumoniae* infection.

While butyrate in uninfected macrophages did not modulate the expression of the polarization markers, it restored *C. pneumoniae* induced suppression of the markers (Figure 6). CD64 was most prominently upregulated (RQ = 2.2), while CD80, TLR4, CD204, and CD206 showed more moderate but clear increase. These results suggest induction of a polyactivated or hybrid macrophage phenotype and align with prior reports describing context-dependent reprogramming of macrophage polarization by butyrate ^28, 57^.

Mechanistically, butyrate’s effects on gene expression are known to involve HDAC inhibition leading to altered chromatin accessibility ^30, 58^. Through HDAC inhibition, butyrate increases transcription which can alter both proinflammatory and anti-inflammatory programs and promote macrophage activation and thus *Chlamydia* clearance. Our data indicate that butyrate may exert transcriptional reprogramming in THP-1 macrophages infected with *C. pneumoniae*.

In fact, previous research connects *Chlamydia* infection with various epigenetic changes driving pathophysiological processes in the host ^59^. One reported example on such chlamydia-host interaction is the ability of *C. trachomatis* effector NUE and its *C. pneumoniae* homologue CpnSET to enter the host cell nucleus and induce histone modifications ^60^. Hayward et al (2020) ^61^ report, that *C. trachomatis* infection induces chromatin loosening and increased transcriptional availability in the HDAC2 and HDAC9 locus in epithelial cells. Even though *Chlamydia*-induced HDAC upregulation has not been explicitly reported in scientific literature, it is possible that in infected THP-1 macrophages, *Chlamydia* effectors upregulate HDAC expression. This could facilitate its own survival by suppressing general transcription in the host cells, which results in decreased macrophage polarization among other actions. Following this logic, butyrate-mediated HDAC inhibition leads to enhanced transcription and macrophage activation, which in turn limits *Chlamydia* survival.

Similar to other HDAC inhibitors, several earlier studies have reported that butyrate upregulates also human antimicrobial peptide, cathelicidin LL-37 expression in various cell lines ^62–64^. In addition to its direct bactericidal properties, LL-37 has several immunomodulatory functions ^65^ and earlier research indicates that some intracellular pathogens are able to suppress LL-37 expression to enhance their survival ^66, 67^. However, the role of LL-37 in chlamydia-host interplay remains unknown. According to our data, *C. pneumoniae* downregulates the LL-37 expression in THP-1 cells (Fig. 7A). This finding aligns with earlier studies showing that *C. trachomatis* can inhibit LL-37 function through plasmid encoded virulence factor Pgp3 ^68, 69^ and chlamydial HtrA protease ^70^.

LL-37 has antibacterial activity against several Gram-positive and Gram-negative bacteria ^42^. It is able to penetrate through cell membranes and thus interact also with intracellular bacteria ^71^. However, previous antichlamydial studies on LL-37 have targeted only the extracellular EB forms of the bacterium. According to these studies, LL-37 can decrease *C. trachomatis* EB infectivity when EBs are pretreated with LL-37 before infecting the host cells ^68, 70^. However, similar experimental procedure on *C. pneumoniae* EBs did not show decreased infectivity by LL-37 treatment ^72^ ^73^, suggesting species-specific differences in EB susceptibility. In addition, earlier research has no evidence on whether LL-37 could inhibit intracellular growth of *Chlamydia*.

To elaborate this question, we conducted a follow up experiment to evaluate the impact of exogenous LL-37 on *C. pneumoniae* genome copy numbers in THP-1 macrophages.

According to our data, LL-37 did not decrease bacterial load in the cells. LL-37 seems to even increase *C. pneumoniae* genome copy numbers compared to the nontreated control (Fig. 7B). One possible explanation to this observation could be LL-37 -induced NLRP3 inflammasome activation. LL-37 expression has been shown to induce caspase-1 and IL-1β expression, triggering NLRP3 inflammasome activation ^74, 75^, which has been reported to promote *C. pneumoniae* replication in cultured macrophages ^76^. Regardless of the underlying molecular mechanisms, our results suggest that LL-37 does not affect intracellular growth of *Chlamydia*, at least in THP-1 macrophages.

Consistent with the above-suggested HDAC-mediated mechanism, Butyrate treatment restored and induced LL-37 expression in *C. pneumoniae* infected cells. Lack of direct effect of LL-37 on chlamydial genome copy numbers suggests that LL-37 induction does not constitute the primary mechanism for butyrate’s antichlamydial effect. Previous research suggests that HDAC inhibition may contribute to enhanced innate defense mechanisms against *Mycobacterium* ^77^, warranting more detailed exploration of this topic also in the context of chlamydial infections.

To conclude, we herein demonstrate that butyrate effectively inhibits *Chlamydia* infection in macrophages, while having no significant impact in epithelial cells. This differential effect may be linked to butyrate’s influence on host immune function, suggesting a potential mechanism for its antichlamydial activity. These results suggest that modulation of host immune function through dietary or microbial metabolites may contribute to the physiological control of chlamydial infections.

## Author contributions

**MY:** Writing – original draft, Visualization, Methodology, Investigation, Formal analysis, Conceptualization, Supervision. **IR:** Writing – review & editing, Methodology, Investigation, Formal analysis, Conceptualization, Supervision. **LH:** Writing – review & editing, Supervision, Project administration, Funding acquisition, Conceptualization.

## Data availability

The data supporting this article have been included as part of the Supplementary Information.

## Acknowledgements

We would like to acknowledge Nana Snellman and Alejandro Cebollada for their contributions to experimental work. This work has been financially supported by Research Council of Finland, grants 333291 and 358425 to LH.

## Abbreviations

AB: aberrant body
BMDM: Bone Marrow-Derived Macrophage
BSA: bovine serum albumin
CD: cluster of differentiation
CHX: cycloheximide
COPD: Chronic obstructive pulmonary disease
EB: elementary body
GAPDH: Glyceraldehyde 3-phosphate dehydrogenase
LPS: lipopolysaccharide
MOI: multiplicity of infection
NLRP3: NOD-LRR- and pyrin domain-containing protein 3
PMA: Phorbol 12-myristate 13-acetate
RT: reticulate body
SCFA: short chain fatty acid
TLR4: Toll-like receptor 4

## Supporting information description

Information of impact of SCFAs on cell viability (Supplementary information 1, PDF). Primers used in the RT-PCR assays (Supplementary information 2, PDF).

## Graphic for table of contents

Note: Created in BioRender. https://BioRender.com

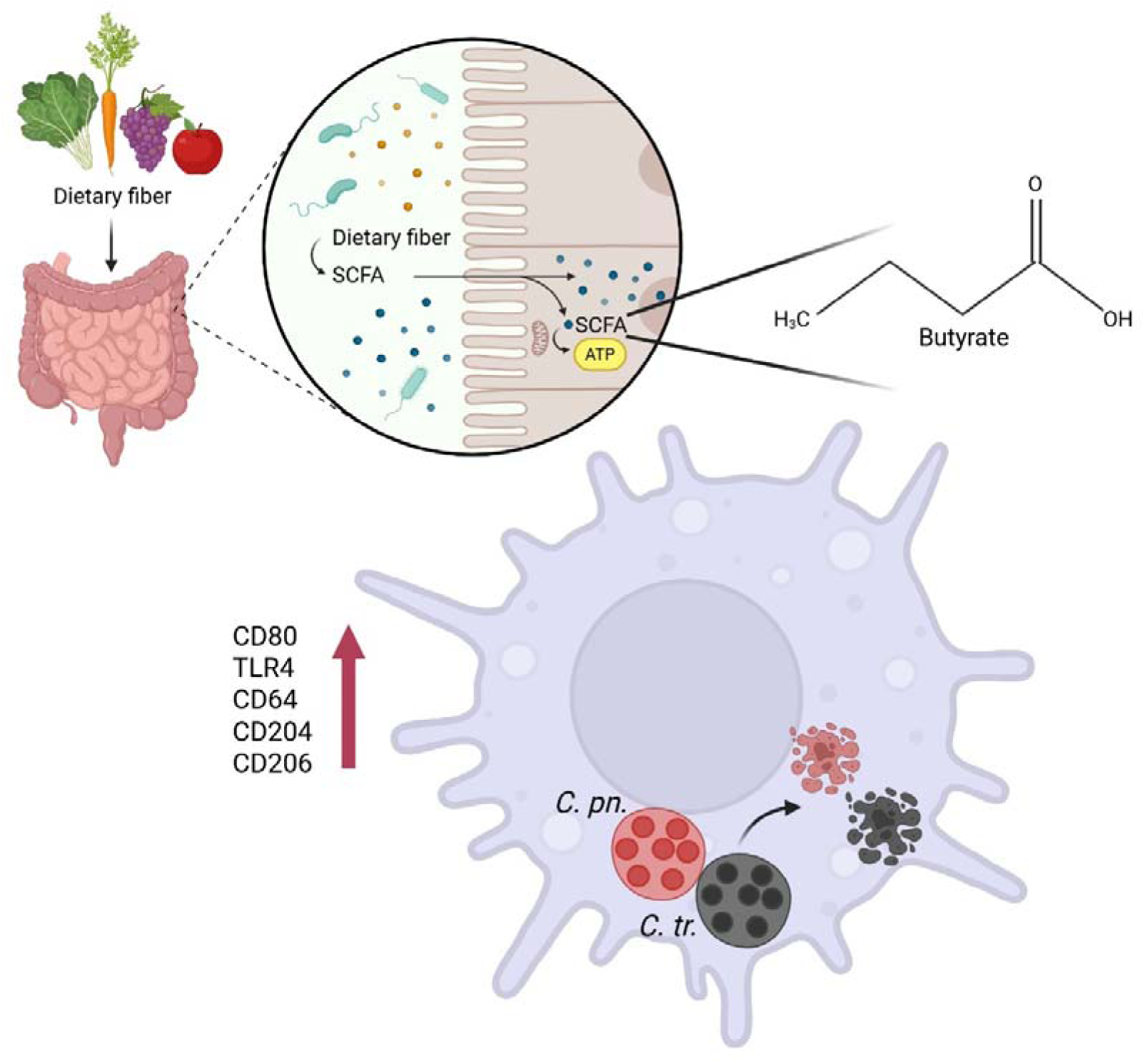

