## Supplementary file 1 for "Butyrate reverts *Chlamydia pneumoniae* -induced transcriptional suppression of macrophage polarization markers and promotes chlamydial clearance in macrophages"

**Supplementary information 1**

**Viability assays**

Viability of A549, HeLa-229 and THP-1 macrophages upon exposure to SCFAs were assessed with resazurin assay. A549 and HeLa cells were seeded into 96-well plate at a density of 6×10^4^ cells/well and incubated overnight. THP-1 cells were seeded into 96-well plate at a density of 8×10^4^ cells/well, exposed to 160 nM PMA and incubated 72 h. Cells were exposed to several concentrations (1000, 700, 500, 400, 300, 200, 100 and 50 µM) of SCFAs and incubated for 48 (HeLa) or 72 h (A549 and THP-1). At the end of the incubation time, samples were removed from the wells and 20 µM resazurin solution was added to the cells and after 2 h incubation, fluorescence was recorded at 570/590 nm with Varioskan Lux plate reader (Thermo Fisher Scientific).

**Impact of SCFA on cell viability**

As shown in Supplementary figure 1, not any of the SCFAs have any effect on the cell viability of either cell line.

**A**

A549

|  | NaAc | NaPr | NaBu |
| --- | --- | --- | --- |
| 1000 μM | 99% | 98% | 91% |
| 700 μM | 97% | 94% | 93% |
| 500 μM | 93% | 94% | 88% |
| 400 μM | 92% | 99% | 85% |
| 300 μM | 91% | 92% | 89% |
| 200 μM | 86% | 89% | 93% |
| 100 μM | 82% | 90% | 95% |
| 50 μM | 88% | 93% | 97% |

**B**

THP-1

|  | NaAc | NaPr | NaBu |
| --- | --- | --- | --- |
| 1000 μM | 104% | 90% | 93% |
| 700 μM | 102% | 101% | 96% |
| 500 μM | 107% | 97% | 97% |
| 400 μM | 91% | 94% | 96% |
| 300 μM | 92% | 95% | 96% |
| 200 μM | 87% | 94% | 92% |
| 100 μM | 96% | 93% | 96% |
| 50 μM | 94% | 98% | 95% |

**C**

HeLa

|  |  |
| --- | --- |
| 1000 μM | 101.216% |
| 700 μM | 96.4035% |
| 500 μM | 95.3704% |
| 400 μM | 90.3145% |
| 300 μM | 84.6065% |
| 200 μM | 84.583% |
| 100 μM | 86.0836% |

Supplementary Figure 1. Impact of SCFA on cell viability of A) A549, B) THP-1 and C) only butyrate on HeLa cells. Results are normalized to nontreated control and presented as mean ± SEM. Statistical significances were calculated using ANOVA and Games-Howell post hoc test.
